# Increased CA3 burst activity in *Doc2α* and *Syt7* knockout mice

**DOI:** 10.64898/2026.07.01.735713

**Authors:** Raghava Jagadeesh Salaka, Edwin R. Chapman

## Abstract

The hippocampal CA3 subfield is central to associative learning and memory consolidation. CA3 pyramidal neurons execute these functions by generating hypersynchronous bursts that feed forward to the CA1. Extensive recurrent collateral connections within CA3 are crucial for the generation of this burst activity. Double C2 domain-containing protein α (Doc2α) and synaptotagmin 7 (Syt7) are high-affinity calcium sensors implicated in asynchronous release (AR) of neurotransmitter and in the exocytosis of dense-core vesicles (DCVs). Additionally, Doc2α is a sensor for miniature neurotransmission, whereas Syt7 is involved in synaptic facilitation and SV replenishment. Both Doc2α and Syt7 are expressed in the hippocampus, but their potential roles in spontaneously bursting networks, like CA3, remain unanswered. Using whole-cell recordings at near physiological conditions (33-34 °C and 1.2 mM Ca^2+^) in acute hippocampal slices obtained from juvenile (P15-21) KO mice lacking Doc2α, Syt7, or both proteins, we report no changes in spontaneous inhibitory or excitatory postsynaptic current frequency or amplitude in CA1 pyramidal neurons. However, there was a marked increase in burst activity, originating in CA3, in all three KO conditions; this was not due to changes in AR or the intrinsic properties of CA3 pyramidal neurons. We propose that this novel burst phenotype in Doc2α- and Syt7-KO mice is unrelated to changes in neurotransmitter release but might be mediated by changes in neuropeptide release from DCVs in the CA3 subfield, or along upstream pathways. Regardless of the underlying mechanisms, this work reveals that both Doc2α and Syt7 act to regulate network activity.

**KEY FINDINGS:**

- Hippocampal CA3 burst activity is critical for associative learning and memory consolidation
- Doc2α and Syt7 are high-affinity Ca^2+^ sensors with similar kinetic properties and are expressed in the hippocampus
- Doc2α and Syt7 support various modes of neurotransmitter release and presynaptic plasticity
- Both Doc2α and Syt7 are also involved in the exocytosis of dense-core vesicles in neuroendocrine cells
- Loss of Doc2α and Syt7 results in increased CA3 burst activity without changing AR or the intrinsic properties of CA3 neurons
- Our results suggest a novel role for Doc2α and Syt7 in regulating CA3 burst activity

**GRAPHICAL ABSTRACT:** The hippocampal CA3 subfield forms an extensive recurrent collateral network enabling the generation of hypersynchronous bursts. In knockout (KO) mice lacking Doc2α, Syt7, or both proteins, there is an increase in CA3 burst activity. In this study, we hypothesize that this increase is likely due to alterations in dense-core vesicle-mediated control of burst activity.

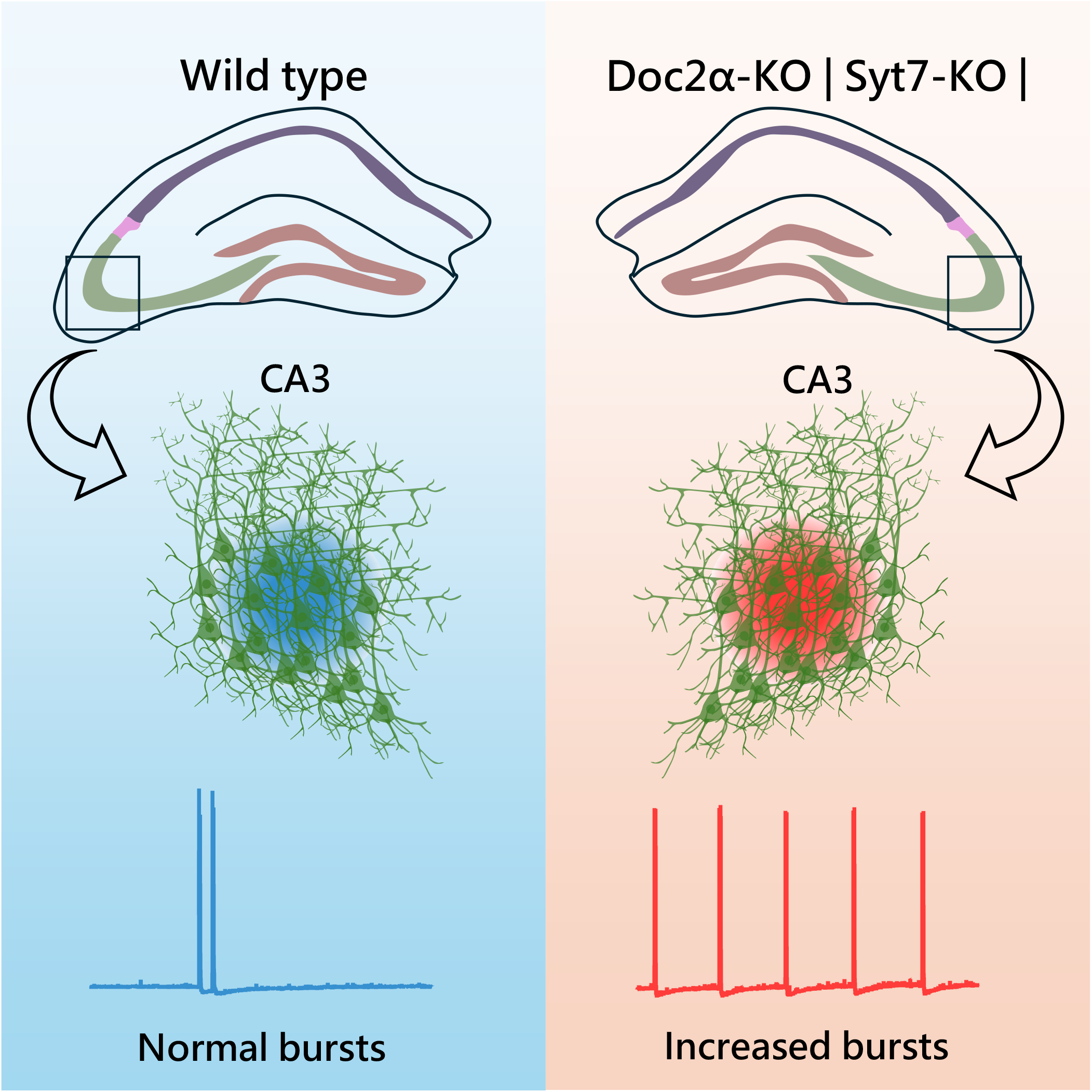

## INTRODUCTION

The hippocampal *cornu ammonis* 3 (CA3) region is critical for associative learning and memory. CA3 pyramidal neurons have axons with extensive recurrent collaterals (RCs) forming synapses within their own population (CA3→CA3) (Rebola *et al*., 2017; Joo & Frank, 2018). Under physiological conditions, CA3 RCs generate hypersynchronous network activity, which causes oscillations in the CA1 subfield via the Schaffer collateral pathway (SC→CA1). This is a crucial process for reliable information transfer and memory consolidation (Buzsáki, 2015). However, increased CA3 burst activity is also a major pathological finding in many neurological disorders, including epilepsy, and the CA3 subfield is a major seizure onset zone in temporal lobe epilepsy (Goldberg & Coulter, 2013).

Double C2 domain-containing protein 2*a* (Doc2α) and synaptotagmin 7 (Syt7) are high-affinity calcium (Ca^2+^) sensors with similar, slow, intrinsic kinetics (Hui *et al*., 2005). Both proteins bind Ca^2+^ via their tandem C2 domains, C2A and C2B. Syt7 is a membrane protein (Vevea *et al*., 2021), whereas Doc2α is a cytosolic protein that translocates to the plasma membrane in response to Ca^2+^ influx (Xue *et al*., 2015). In neuroendocrine cells, both proteins serve as Ca^2+^ sensors that trigger dense-core vesicle (DCV) exocytosis and the release of neuropeptides and hormones (MacDougall *et al*., 2018; van Westen *et al*., 2021). In neurons, Doc2α (and a closely related isoform, Doc2β) functions as a Ca^2+^ sensor that regulates spontaneous neurotransmitter release (Groffen *et al*., 2010; Courtney *et al*., 2018), and Syt7 regulates synaptic facilitation and synaptic vesicle (SV) replenishment (Chen *et al*., 2017; Jackman & Regehr, 2017). In addition, Doc2α and Syt7 play a role in the asynchronous release (AR) of neurotransmitters from nerve terminals (Yao *et al*., 2011; Xue *et al*., 2015; Chen *et al*., 2017; Turecek & Regehr, 2018; Deng *et al*., 2020; Wu *et al*., 2024).

Asynchronously released quanta from excitatory terminals would maintain the post-synaptic neuron in a depolarized state for prolonged periods, enhancing the spiking window and temporal precision of AP firing (Iremonger & Bains, 2007; Evstratova *et al*., 2014). Knocking out proteins important for AR would be expected to reduce the excitation of the postsynaptic neuron. Consistent with this hypothesis, in our previous study, knockdown (KD) of *Doc2α* reduced reverberatory activity in primary neuronal cultures (Yao *et al*., 2011). Similarly, in cerebellum, *Syt7* KOs showed a significant reduction in molecular layer interneuron firing after parallel fiber stimulation (Weingarten *et al*., 2024). Also, desynchronization of glutamate release from CA3 terminals by substituting Ca^2+^ with strontium (Sr^2+^) prolongs burst duration in CA3 (Jones *et al*., 2007). So, we hypothesized that loss of Doc2α or Syt7 would decrease bursting activity in CA3, as both proteins are expressed in these cells (Verhage *et al*., 1997; Jackman & Regehr, 2017). We therefore performed whole-cell voltage and current clamp recordings in CA1, to assess CA3 burst firing, in acute hippocampal slices from juvenile mice lacking these proteins. Surprisingly, we observed an increase in CA3 burst generation in *Doc2α-* and *Syt7-*KO mice following pharmacological disinhibition. The increased burst activity does not appear to be due to changes in neurotransmitter release mechanisms or the intrinsic properties of CA3 neurons, indicating a novel role for Doc2α and Syt7 in network activity. We propose that these two Ca^2+^ sensors affect burst generation by regulating the release of neuropeptides in the hippocampus.

## METHODS

### Ethical Approval

Animal experiments were performed in accordance with the National Institutes of Health guidelines for animal research and were approved by the Animal Care and Use Committee (IACUC, assurance number: A3368-01) at the University of Wisconsin-Madison.

### Animals

*Doc2α* knockout (*Doc2α*^−/−^) (Sakaguchi *et al*., 1999), *Syt7* knockout (*Syt7*^−/−^) (Chakrabarti *et al*., 2003) and double knockout (DKO) (*Doc2α*^−/−^/*Syt7*^−/−^) (Wu *et al*., 2024) were used as previously described. Homozygous KOs and WT breeders were derived from heterozygous mice of each strain. Heterozygous mice were backcrossed to C57BL6/J approximately every five generations. Offspring from homozygous breeding pairs were not used to create additional matings. All mice were housed in polycarbonate cages on a 12h dark/light cycle and with food and water provided *ad libitum*. We did not observe any sex differences; therefore, data from both sexes were pooled.

### Acute slice preparation

P15-21 mice were deeply anaesthetized with isoflurane inhalation in a 1 L covered chamber precharged with isoflurane-dabbed tissue paper placed in a mesh metal sphere. Mice were introduced one minute later and observed for decreased breathing frequency and increased gasping. Pain reflexes were assessed by tail pinch, and a lack of response was considered evidence of mouse unconsciousness. Mice were then decapitated, and brains were quickly removed and chilled in ice-cold sucrose-based cutting solution containing (in mM): 210 sucrose, 2.5 KCl, 25 NaHCO_3_, 1.25 NaH_2_PO_4_, 20 glucose, 7 MgSO_4_ 7H_2_O and 0.5 CaCl_2_ 2H_2_O. Two minutes later, hemispheres were isolated, and modified transverse hippocampal slices were cut at 300 μm thickness using the 10° angle blocking cut (Bischofberger *et al*., 2006) with a vibratome (VT1200s, Leica Microsystems, RRID: SCR_018453) in the same solution. Slices were transferred to an incubator (BSK1, Automate Scientific, RRID: SCR_029006) filled with prewarmed artificial cerebrospinal fluid (aCSF) (~34°C) for 20 min and then stored at room temperature until recording (6-8 h). The aCSF contained the following components (in mM): 125 NaCl, 2.5 KCl, 25 NaHCO_3_, 1.25 NaH_2_PO_4_, 20 glucose, 1 MgSO_4_ 7H_2_O, and 1.2 CaCl_2_ 2H_2_O. The osmolarity of the aCSF was measured using a vapor pressure osmometer (VAPRO^®^ Model 5600, Elitech Group Inc.) and was adjusted to 300-305 mmol/kg. Throughout the procedure, slices were bubbled with a 95% O_2_ and 5% CO_2_ mixture (carbogen). Slices were transferred to a submersion chamber (RC26GLP, Warner Instruments Company, RRID: SCR_029015) that is continuously perfused with aCSF at a rate of 2-3ml/min using a peristaltic pump (Miniplus 3, Gilson Inc., RRID: SCR_029008). All the recordings were performed at 33-34° C using an in-line heater (SH-27B, Warner Instrument Company) with a temperature controller (TC-324C, Warner Instrument Company), and the solutions were not recycled. Whole-cell voltage clamp or current clamp recordings were obtained from CA1 and CA3 neurons visualized via a 60X water immersion lens (LUMPLFLN60XW, N. A. = 1) attached to an upright microscope (Olympus BX51WI, Evident Scientific Inc., RRID: SCR_023069) equipped with DIC and a CCD camera (Retiga ELECTRO, Teledyne Technologies Inc.). Neurons were patched using borosilicate glass pipettes (BF150-110-10HP, Sutter Instrument Company) with a pipette resistance (*R*_*p*_) of 2-5 MΩ, pulled using a Flaming/Brown pipette puller (P1000, Sutter Instrument Company, RRID: SCR_021042).

### Whole-cell voltage clamp recordings in CA1

Spontaneous excitatory postsynaptic currents (sEPSCs) were obtained in the presence of 100 μM picrotoxin (Hellobio) at −70 mV for 5 minutes using a pipette internal solution comprising (in mM): 135 cesium-methanesulfonate, 10 HEPES, 8 NaCl, 5 QX-314 (Hellobio), 2 Mg-ATP, 1 EGTA, and 0.3 sodium-GTP hydrate (osmolarity = 290 mmol/kg)(Barragan *et al*., 2024). Spontaneous inhibitory postsynaptic currents (sIPSCs) were obtained in the presence of 10 μM CNQX (Hellobio) and 50 μM APV (Hellobio) at −60 mV for 5 minutes using a pipette internal solution composed of (in mM): 120 cesium chloride, 10 HEPES, 5 phosphocreatine, 5 QX-314 (Hellobio), 2 Mg-ATP, 5 EGTA, and 0.3 sodium-GTP hydrate, 2 MgCl_2_ and 2CaCl_2_ (osmolarity = 290 mmol/kg) (Viana da Silva *et al*., 2019). All chemicals used throughout this study were from Sigma unless otherwise specified. sIPSCs were analyzed for only 1 minute, due to the high event frequency. For asynchronous neurotransmitter release (AR) measurements, a cut was made in the CA3/CA2 border to prevent recurrent activity. A platinum-iridium cluster bipolar electrode (E000010596, FHC Inc.) was placed ~50 µm deep in the *stratum radiatum* of the CA1 region to stimulate the Schaffer collaterals every 8 s (0.125 Hz). Evoked excitatory postsynaptic currents (eEPSCs) were recorded in the presence of 100 μM picrotoxin and 50 μM APV at −70 mV. At least 100 sweeps were recorded and analyzed per neuron. Responses were elicited using 0.2 ms square-wave pulses delivered via a stimulus isolation unit (A385, World Precision Instruments) controlled by software through TTL gating. The stimulus intensity varied across experiments (40-140 μA) and was adjusted to elicit a response of 0.8-1.5 nA. AR events were normalized to the evoked synchronous EPSC amplitude (Weingarten *et al*., 2024). The series resistance (*R*_*s*_) was less than 15 MΩ for AR measurements and 25 MΩ for other recordings. *R*_*s*_ was not compensated and was monitored offline. Recordings were discarded if the *R*_*s*_ changed by 30% during the experiment or if the holding current exceeded −250 pA. All recordings were obtained using an amplifier (Multiclamp 700B, Molecular Devices, RRID: SCR_018455) and a digitizer (Digidata 1550B, Molecular Devices) interfaced with software (Clampex v11.3, pClamp Suite, Molecular Devices, RRID: SCR_011323). The signals were filtered at 2 kHz using a Bessel filter and digitized at 10 kHz.

### Whole-cell current clamp recordings in CA1 and CA3

Two types of current clamp measurements were performed: 1) spontaneous excitatory postsynaptic potentials (sEPSPs) from CA1 neurons, and 2) intrinsic properties of CA3 neurons. Neurons in the CA1 region, or near the CA3a and CA3b border, were patched with a K^+^-containing internal solution that had the following components (in mM): 135 K-gluconate, 10 HEPES, 10 Na-phosphocreatine, 5 NaCl, 4 Mg-ATP, 2 MgCl2, 0.4 Na-GTP and 0.2 EGTA (osmolarity = 290 mmol/kg). The resting membrane potential (RMP, V_*rest*_) was measured immediately after breaking-in; we then allowed 2-3 minutes for dialysis of the internal solution in the patched neurons. For CA1 experiments, sEPSPs were recorded for 5 minutes. CA1 recordings were performed at RMP (*I* = 0 mode) and discarded if the membrane potential was not −60 mV or lower. The bath perfusion aCSF for these experiments contained 100 μM picrotoxin (Hellobio). To assess the intrinsic properties of CA3 neurons, a membrane potential of −70 ± 2 mV was maintained by injecting a positive or negative current of 50 to −150 pA. Recordings were performed with aCSF solution containing 100 μM picrotoxin, 10 μM CNQX disodium salt (Hellobio) and 50 μM DL-APV sodium salt (Hellobio). The firing rate of the CA3 neurons was assessed by injecting a 25 pA depolarizing current step of 500 ms duration in increasing order (i.e., 0, 25, 50, 75, 100, 125, 150, 175, 200, and 0). The number of action potentials was counted at each step. For input resistance, a hyperpolarizing current of 500 ms duration was injected in −20 pA stepwise increments between 0 to −200 pA. The average change in voltage over the last 100 ms was taken as the steady-state response. The slope of a regression line fitted to the voltage-current relationship was used to calculate the overall input resistance. Action potential (AP) height and width were assessed by injecting a single 2 ms 1800 pA current pulse. AP amplitude was calculated as the difference between the baseline and the peak. AP half-width was measured at 50% of the peak-to-baseline amplitude. All current clamp protocols were repeated at least 3 times, and the data were averaged. Junction potentials were not corrected. The signals were filtered at 2 kHz using a Bessel filter and digitized at 10 or 20 kHz.

### Data Analysis

sEPSCs, sIPSCs and quantal AR events were detected using a template-matching algorithm in Easy Electrophysiology® (Easy Electrophysiology Ltd, RRID: SCR_021190). The detection limit was set to 3 times the root-mean-square of the peak-to-peak noise, which was typically around 4-6 pA for EPSCs and 8-10 pA for IPSCs. Burst detection was performed manually with Clampfit v11.3 software. In voltage clamp mode, large-amplitude (>200 pA) and long-duration (>100 ms) currents were defined as burst-like events and their duration was excluded when calculating the sEPSC frequency. In current clamp mode, bursts were characterized by 3-22 APs throughout the burst, or by 3-4 APs during the initial phase followed by a prolonged depolarization in the middle, and 1-2 APs at the end. Burst duration was determined as the interval between the very start of the depolarization from the baseline to the initial return to baseline during repolarization phase. Intrinsic properties were analyzed using Clampfit v11.3 (pClamp Suite, Molecular Devices Inc., RRID: SCR_011323).

### Statistical Analysis

All data are presented as mean ± SD, with *n* = number of neurons. Statistical analysis and graphs were generated using Prism v11 (GraphPad Inc., RRID: SCR_002798). Data were tested for normality (Shapiro-Wilk test), and based on the distribution, either the Brown-Forsythe ANOVA test or the Kruskal-Wallis test was used with a multiple comparisons test for post-hoc correction. A *p-*value less than 0.05 was considered significant. Additional details regarding the statistical methods are presented in the Results and Figure Legends.

## RESULTS

### sIPSC and sEPSC frequency is unaltered in *Doc2α*- and *Syt7*-KOs

Given the numerous roles of Doc2α and Syt7 in the exocytosis of SVs and DCVs (Friedrich *et al*., 2008; Ramalingam *et al*., 2012; Houy *et al*., 2017; MacDougall *et al*., 2018; van Westen *et al*., 2021), we tested whether loss of these proteins affects network-dependent spontaneous activity in the mouse hippocampus, to alter the excitatory-inhibitory balance. We carried out whole-cell voltage-clamp recordings in CA1 pyramidal neurons in acute slice preparations. We first tested whether the inhibitory network in this region is affected in the KOs by recording sIPSCs. CA1 cells receive numerous inhibitory inputs from different layers of the hippocampus (Fig. 1A) (Megías *et al*., 2001). We found no significant changes in sIPSC frequency (Fig. 1B and C; *p* = 0.148, Brown-Forsythe ANOVA test with Dunnett T3 multiple comparisons test; WT = 13.47 ± 6.73 Hz; *Doc2α*^−/−^ = 16.52 ± 7.20; *Syt7*^−/−^ = 18.29 ± 7.69 Hz; *Doc2α*^−/−^*/Syt7*^−/−^ = 18.66 ± 7.69 Hz; WT = 20 cells from 11 slices from 6 mice; *Doc2α*^−/−^ = 21 cells from 12 slices from 7 mice; *Syt7*^−/−^ = 16 cells from 9 slices from 5 mice; *Doc2α*^−/−^*/Syt7*^−/−^ = 14 cells from 7 slices from 5 mice) or amplitude (Fig. 1B and D; *p* = 0.520, Kruskal-Wallis test with Dunn’s multiple comparisons test; WT = −43.03 ± 14.56 pA; *Doc2α*^−/−^ = −49.92 ± 18.32 pA; *Syt7*^−/−^ = −46.10 ± 11.31 pA; *Doc2α*^−/−^*/Syt7*^−/−^ = −48.30 ± 13.08 pA; WT = 20 cells from 11 slices from 6 mice; *Doc2α*^−/−^ = 21 cells from 12 slices from 7 mice; *Syt7*^−/−^ = 16 cells from 9 slices from 5 mice; *Doc2α*^−/−^*/Syt7*^−/−^ = 14 cells from 7 slices from 5 mice).

**Figure 1.**
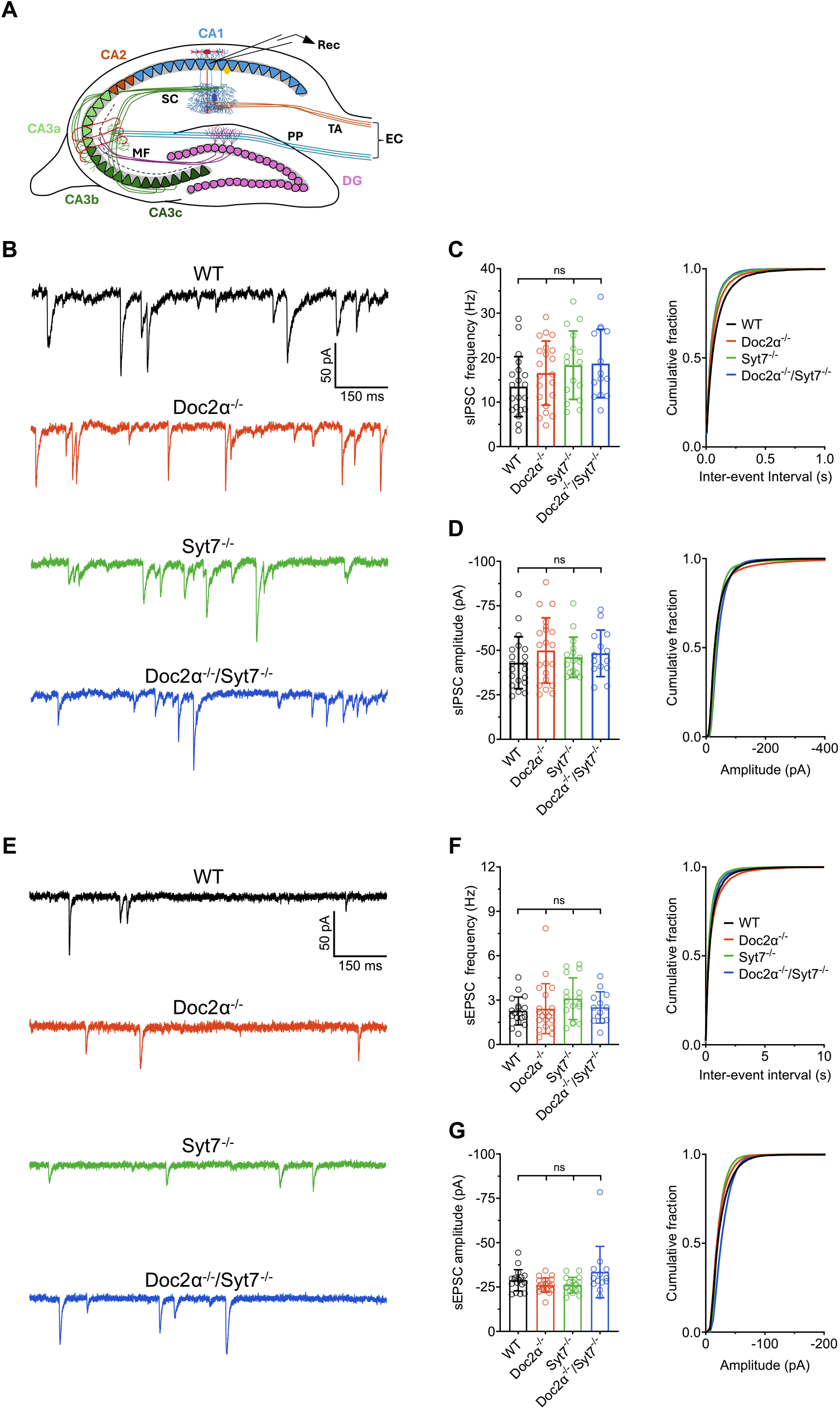
Network-dependent inhibitory and excitatory spontaneous activity is unchanged in *Doc2α*^−/−^, *Syt7*^−/−^ and *Doc2α*^−/−^*/Syt7*^−/−^ mice. A: Schematic diagram of the hippocampal slice preparation showing the inhibitory and excitatory inputs to CA1 pyramidal neurons and whole-cell recording electrode placement. B: Representative sIPSC traces of recordings obtained from CA1 pyramidal neurons. C: sIPSC frequency (left panel) and cumulative distribution (right panel). No significant change in sIPSC frequency was observed between any of the groups (WT = 13.47 ± 6.73 Hz; *Doc2α*^−/−^ = 16.52 ± 7.20; *Syt7*^−/−^ = 18.29 ± 7.69 Hz; *Doc2α*^−/−^*/Syt7*^−/−^ = 18.66 ± 7.69 Hz). WT vs. *Doc2α*^−/−^: *p* = 0.656; WT vs. *Syt7*^−/−^: *p* = 0.287; WT vs. *Doc2α*^−/−^*/Syt7*^−/−^: *p* = 0.261; *Doc2α*^−/−^ vs. *Syt7*^−/−^: *p* = 0.976; *Doc2α*^−/−^ vs. *Doc2α*^−/−^*/Syt7*^−/−^: *p* = 0.9528; *Syt7*^−/−^ vs. *Doc2α*^−/−^*/Syt7*^−/−^: *p* = 0.999. D: sIPSC amplitude (left panel) and cumulative distribution (right panel). No significant change in sIPSC amplitude was observed between any of the groups (WT = −43.03 ± 14.56 pA; *Doc2α*^−/−^ = −49.92 ± 18.32 pA; *Syt7*^−/−^ = −46.10 ± 11.31 pA; *Doc2α*^−/−^*/Syt7*^−/−^ = −48.30 ± 13.08 pA). WT vs. *Doc2α*^−/−^: *p* = 0.999; WT vs. *Syt7*^−/−^: *p* = 0.999; WT vs. *Doc2α*^−/−^*/Syt7*^−/−^: *p* = 0.999; *Doc2α*^−/−^ vs. *Syt7*^−/−^: *p* = 0.999; *Doc2α*^−/−^ vs. *Doc2α*^−/−^*/Syt7*^−/−^: *p* = 0.999; *Syt7*^−/−^ vs. *Doc2α*^−/−^*/Syt7*^−/−^: *p* = 0.999. E: Representative sEPSC traces of recordings obtained from CA1 pyramidal neurons. F: sEPSC frequency (left panel) and cumulative distribution (right panel). No significant change in sEPSC frequency was observed between any of the groups (WT = 2.26 ± 0.94 Hz; *Doc2α*^−/−^ = 2.41 ± 1.68 Hz; *Syt7*^−/−^ = 3.09 ± 1.40 Hz; *Doc2α*^−/−^*/Syt7*^−/−^ = 2.50 ± 1.05 Hz). WT vs. *Doc2α*^−/−^: *p* = 0.999; WT vs. *Syt7*^−/−^: *p* = 0.421; WT vs. *Doc2α*^−/−^*/Syt7*^−/−^: *p* = 0.999; *Doc2α*^−/−^ vs. *Syt7*^−/−^: *p* = 0.197; *Doc2α*^−/−^ vs. *Doc2α*^−/−^*/Syt7*^−/−^: *p* = 0.999; *Syt7*^−/−^ vs. *Doc2α*^−/−^*/Syt7*^−/−^: *p* = 0.999. G: sEPSC amplitude (left panel) and cumulative distribution (right panel). No significant change in sEPSC amplitude was observed between any of the groups (WT = −28.82 ± 5.99; *Doc2α*^−/−^ = −26.04 ± 4.02 pA; *Syt7*^−/−^ = −26.09 ± 4.42 pA; *Doc2α*^−/−^*/Syt7*^−/−^ = −33.44 ± 14.46 pA). WT vs. *Doc2α*^−/−^: *p* = 0.836; WT vs. *Syt7*^−/−^: *p* = 0.916; WT vs. *Doc2α*^−/−^*/Syt7*^−/−^: *p* = 0.999; *Doc2α*^−/−^ vs. *Syt7*^−/−^: *p* = 0.999; *Doc2α*^−/−^ vs. *Doc2α*^−/−^*/Syt7*^−/−^: *p* = 0.140; *Syt7*^−/−^ vs. *Doc2α*^−/−^*/Syt7*^−/−^: *p* = 0.166. In panel C, data were analyzed using the Brown-Forsythe ANOVA test with Dunnett T3 multiple comparisons test. In panels D, F and G, data were analyzed using the Kruskal-Wallis test with Dunn’s multiple comparisons test. For sIPSC frequency and amplitude, WT = 20 cells from 11 slices from 6 mice; *Doc2α*^−/−^ = 21 cells from 12 slices from 7 mice; *Syt7*^−/−^ = 16 cells from 9 slices from 5 mice; *Doc2α*^−/−^*/Syt7*^−/−^ = 14 cells from 7 slices from 5 mice. For sEPSC frequency and amplitude, WT = 18 cells from 10 slices from 6 mice; *Doc2α*^−/−^ = 20 cells from 10 slices from 6 mice; *Syt7*^−/−^ = 17 cells from 8 slices from 6 mice; *Doc2α*^−/−^*/Syt7*^−/−^ = 13 cells from 6 slices from 4 mice. For cumulative distribution plots, only mean values were presented for clarity. Graphs represent mean ± SD. ns: no significance; CA: cornu ammonis; DG: dentate gyrus; EC: entorhinal cortex; PP: perforant pathway; MF: mossy fiber pathway; SC: Schaffer-collateral pathway; TA: temporoammonic pathway; Rec: recording electrode.

We next explored potential changes in network-dependent spontaneous excitatory activity. The CA1 region receives extensive excitatory inputs from CA3 neurons via Schaffer collaterals and from layer-III pyramidal neurons of the entorhinal cortex via the temporoammonic pathway (Fig. 1A). Analogous to our findings regarding sIPSCs, we observed no significant changes in sEPSC frequency (Fig. 1E and F; *p* = 0.192; Kruskal-Wallis test with Dunn’s multiple comparisons test; WT = 2.26 ± 0.94 Hz; *Doc2α*^−/−^ = 2.41 ± 1.68 Hz; *Syt7*^−/−^ = 3.09 ± 1.40 Hz; *Doc2α*^−/−^*/Syt7*^−/−^ = 2.50 ± 1.05 Hz; WT = 18 cells from 10 slices from 6 mice; *Doc2α*^−/−^ = 20 cells from 10 slices from 6 mice; *Syt7*^−/−^ = 17 cells from 8 slices from 6 mice; *Doc2α*^−/−^*/Syt7*^−/−^ = 13 cells from 6 slices from 4 mice) or amplitude (Fig. 1E and G; *p* = 0.636; Kruskal-Wallis test with Dunn’s multiple comparisons test; WT = −28.82 ± 5.99; *Doc2α*^−/−^ = −26.04 ± 4.02 pA; *Syt7*^−/−^ = −26.09 ± 4.42 pA; *Doc2α*^−/−^*/Syt7*^−/−^ = −33.44 ± 14.46 pA; WT = 18 cells from 10 slices from 6 mice; *Doc2α*^−/−^ = 20 cells from 10 slices from 6 mice; *Syt7*^−/−^ = 17 cells from 8 slices from 6 mice; *Doc2α*^−/−^*/Syt7*^−/−^ = 13 cells from 6 slices from 4 mice).

### Loss of either Doc2α or Syt7 increases CA3 burst generation

We next turned our attention to CA3 burst generation. The activity of a single CA3 neuron is sufficient to trigger a burst within the subfield when GABAergic activity is inhibited with antagonists. For example, 100 µM picrotoxin is widely used to study CA3 burst generation and propagation (Miles & Wong, 1983; Hablitz, 1984; Yee *et al*., 2003; Cohen *et al*., 2006; Jones *et al*., 2007; Shao & Dudek, 2009; Buzsáki, 2015). Under these experimental conditions, hypersynchronous bursts are generated due to spontaneous activity and an extensive RC network in the CA3 subfield, and then spread to the CA1 subfield via a feedforward mechanism (Shao & Dudek, 2009). In recordings of CA1 spontaneous excitatory activity, we observed an increase in the number of burst-like events with large amplitude (>200 pA) and long duration (>100 ms) currents; this increase was 2-4-fold in each individual KO as well as the DKO condition compared to WT, but not between KO groups (Fig. 2A and B; *p* < 0.0001; Kruskal-Wallis test with Dunn’s multiple comparisons test; WT = 2.44 ± 2.41 burst-like events/min; *Doc2α*^−/−^ = 4.98 ± 2.55 burst-like events/min; *Syt7*^−/−^ = 9.59 ± 4.65 burst-like events/min; *Doc2α*^−/−^*/Syt7*^−/−^ = 6.86 ± 4.31 burst-like events/min; WT = 18 cells from 10 slices from 6 mice; *Doc2α*^−/−^ = 20 cells from 10 slices from 6 mice; *Syt7*^−/−^ = 17 cells from 8 slices from 6 mice; *Doc2α*^−/−^*/Syt7*^−/−^ = 13 cells from 6 slices from 4 mice). Moreover, the burst-like events were absent when a cut was made (see Fig. 4A for an illustration of the cut angle) across the Schaffer collateral projections between the CA3 and CA1 (Fig. 2C; WT = 14 cells from 5 slices from 4 mice; *Doc2α*^−/−^ = 12 cells from 5 slices from 4 mice; *Syt7*^−/−^ = 16 cells from 7 slices from 4 mice; *Doc2α*^−/−^*/Syt7*^−/−^ = 12 cells from 6 slices from 5 mice.)

**Figure 2.**
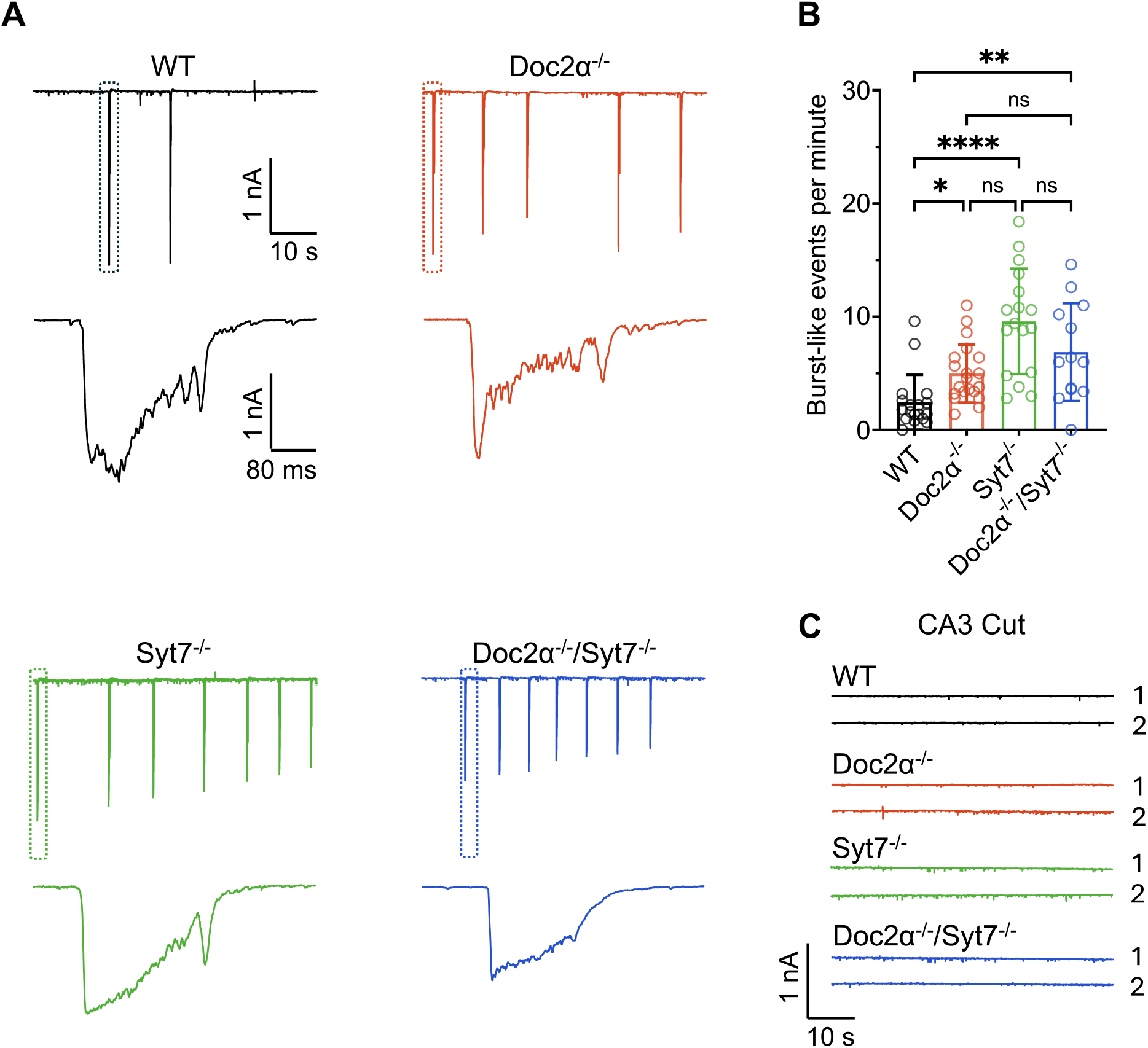
Increased spontaneous burst-like events at the Schaffer collateral-CA1 synapses in voltage clamp recordings. A: Representative sEPSC traces showing burst-like events (top panel); the area indicated by the dotted rectangle is expanded along the time axis (lower panel) to show the structure of a burst-like event. B: Significant increase in the number of burst-like events per minute in *Doc2α*^−/−^, *Syt7*^−/−^, and *Doc2α*^−/−^*/Syt7*^−/−^ mice (WT = 2.44 ± 2.41 burst-like events/min; *Doc2α*^−/−^ = 4.98 ± 2.55 burst-like events/min; *Syt7*^−/−^ = 9.59 ± 4.65 burst-like events/min; *Doc2α*^−/−^*/Syt7*^−/−^ = 6.86 ± 4.31 burst-like events/min). WT vs. *Doc2α*^−/−^: \**p* = 0.023; WT vs. *Syt7*^−/−^: \*\*\*\**p* < 0.0001; WT vs. *Doc2α*^−/−^*/Syt7*^−/−^: \*\**p* = 0.002; *Doc2α*^−/−^ vs. *Syt7*^−/−^: *p* = 0.078; *Doc2α*^−/−^ vs. *Doc2α*^−/−^*/Syt7*^−/−^: *p* = 0.999; *Syt7*^−/−^ vs. *Doc2α*^−/−^*/Syt7*^−/−^: *p* = 0.999. C: Representative sEPSC traces of recordings without any burst-like events when CA3 was cut; data could not be quantified as no bursts were detected In panel B, data were analyzed using the Kruskal-Wallis test with Dunn’s multiple comparisons test. For bursts per minute data, WT = 18 cells from 10 slices from 6 mice; *Doc2α*^−/−^ = 20 cells from 10 slices from 6 mice; *Syt7*^−/−^ = 17 cells from 8 slices from 6 mice; *Doc2α*^−/−^*/Syt7*^−/−^ = 13 cells from 6 slices from 4 mice. For CA3 cut experiments, WT = 14 cells from 5 slices from 4 mice; *Doc2α*^−/−^ = 12 cells from 5 slices from 4 mice; *Syt7*^−/−^ = 16 cells from 7 slices from 4 mice; *Doc2α*^−/−^*/Syt7*^−/−^ = 12 cells from 6 slices from 5 mice. Graphs represent mean ± SD. ns: no significance.

To better understand the burst-like events and their duration, we patched CA1 pyramidal neurons under the same experimental conditions, but in current-clamp mode from a separate cohort of animals. We again observed a 2-4 fold increase in burst frequency in *Doc2α* KO, *Syt7* KO, and DKO mice compared to WT, but not between KO groups (Fig. 3A and B; *p* < 0.0001; Kruskal-Wallis test with Dunn’s multiple comparisons test; WT = 1.82 ± 1.68 bursts/min; *Doc2α*^−/−^ = 5.62 ± 3.16 bursts/min; *Syt7*^−/−^ = 6.91 ± 4.46 bursts/min; *Doc2α*^−/−^*/Syt7*^−/−^ = 5.20 ± 2.89 bursts/min; WT = 17 cells from 9 slices from 6 mice; *Doc2α*^−/−^ = 17 cells from 11 slices from 7 mice; *Syt7*^−/−^ = 17 cells from 9 slices from 5 mice; *Doc2α*^−/−^*/Syt7*^−/−^ = 18 cells from 8 slices from 4 mice). The number of APs per burst was slightly decreased in the *Syt7* KO compared to the WT, but not in *Doc2α-*KO and DKO (Fig. 3C; *p* < 0.0001, Kruskal-Wallis test with Dunn’s multiple comparisons test; WT = 5.71 ± 2.40 APs/burst; *Doc2α*^−/−^ = 7.76 ± 4.72 APs/burst; *Syt7*^−/−^ = 5.27 ± 3.75 APs/burst; *Doc2α*^−/−^*/Syt7*^−/−^ = 5.36 ± 3.59 APs/burst, 3-5 bursts analyzed per cell; WT = 62 bursts; *Doc2α*^−/−^ = 85 bursts; *Syt7*^−/−^ = 84 bursts; *Doc2α*^−/−^*/Syt7*^−/−^ = 90 bursts). Further analysis revealed that the burst duration was unaltered in the KOs and the DKO compared to WT (Fig. 3D; *p* < 0.0001, Kruskal-Wallis test with Dunn’s multiple comparisons test; WT = 268.40 ± 87.42 ms; *Doc2α*^−/−^ = 307.40 ± 75.05 ms; *Syt7*^−/−^ = 244.70 ± 98.97 ms; *Doc2α*^−/−^*/Syt7*^−/−^ = 246.80 ± 99.37 ms; WT = 62 bursts; *Doc2α*^−/−^ = 85 bursts; *Syt7*^−/−^ = 84 bursts; *Doc2α*^−/−^*/Syt7*^−/−^ = 90 bursts). The bursts were characterized by 3-22 APs throughout the burst, or by 3-4 APs during the initial phase followed by a prolonged depolarization in the middle, and 1-2 APs at the end. Taken together, these findings indicate that bursts originated in CA3 and were driven more strongly in both KOs. It is unclear whether these *in vitro* bursts correspond to enhanced *in vivo* sharp wave (SPW) generation or represent pathological epileptiform activity, but we note that these KO mice do not exhibit overt seizures.

**Figure 3.**
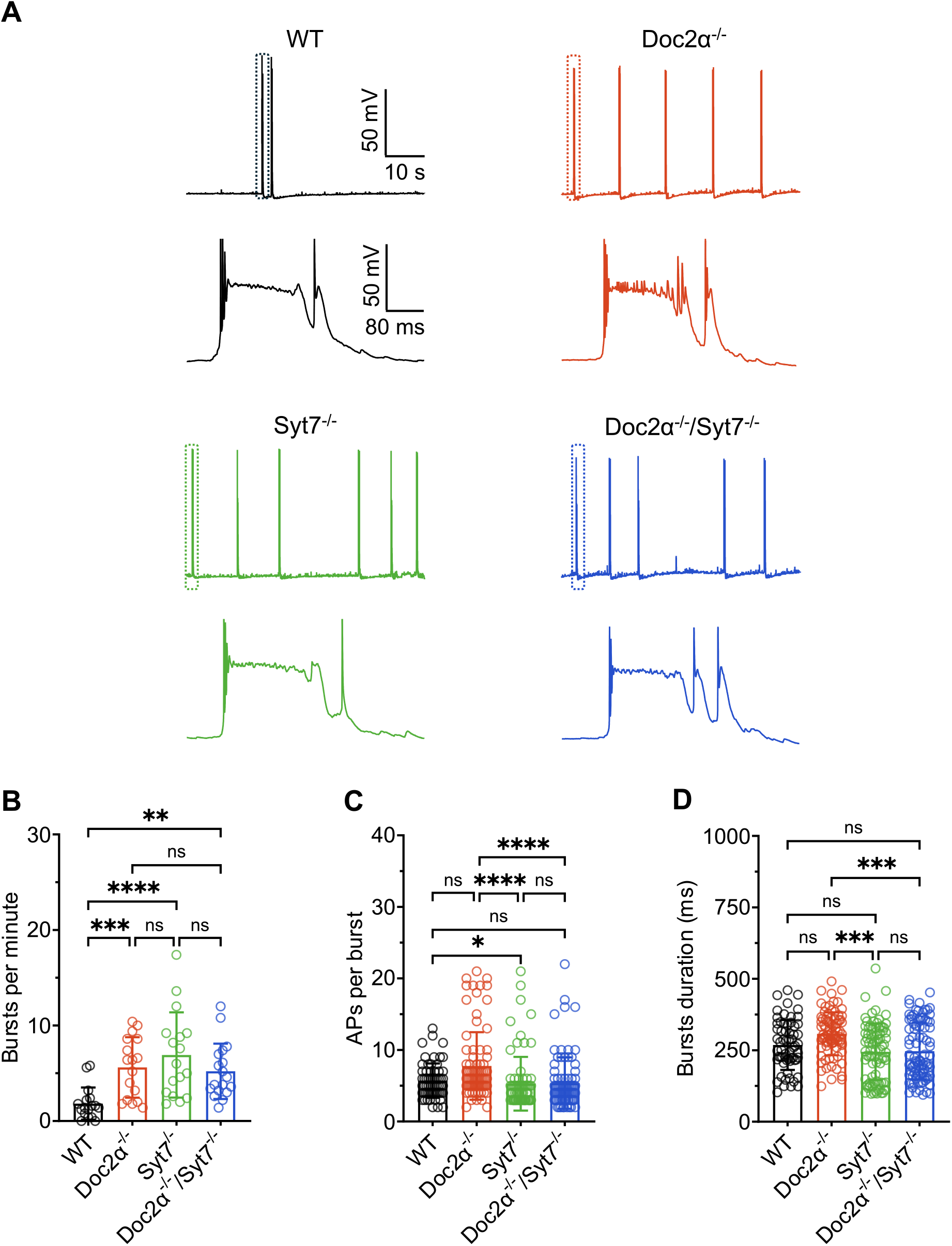
Increased spontaneous bursts at the Schaffer collateral-CA1 synapses in current clamp recordings. A: Representative sEPSP traces showing bursts (top panel); the area indicated by the dotted rectangle is expanded along the time axis (lower panel) to show the structure of a burst. B: Significant increase in the number of bursts per minute in *Doc2α*^−/−^, *Syt7*^*-/*^ and *Doc2α*^−/−^*/Syt7*^−/−^ mice (WT = 1.82 ± 1.68 bursts/min; *Doc2α*^−/−^ = 5.62 ± 3.16 bursts/min; *Syt7*^−/−^ = 6.91 ± 4.46 bursts/min; Doc2α^−/−^/Syt7^−/−^ = 5.20 ± 2.89 bursts/min). WT vs. *Doc2α*^−/−^: \*\*\**p* = 0.001; WT vs. *Syt7*^−/−^: \*\*\*\**p* = 0.0001; WT vs. *Doc2α*^−/−^*/Syt7*^−/−^: \*\**p* = 0.001; *Doc2α*^−/−^ vs. *Syt7*^−/−^: *p* = 0.999; *Doc2α*^−/−^ vs. *Doc2α*^−/−^*/Syt7*^−/−^: *p* = 0.999; *Syt7*^−/−^ vs. *Doc2α*^−/−^*/Syt7*^−/−^: *p* = 0.999. C: Number of action potentials per burst is decreased in *Syt7*^−/−^ but not in *Doc2α*^−/−^ or *Doc2α*^−/−^*/Syt7*^−/−^ mice (WT = 5.71 ± 2.40 APs/burst; *Doc2α*^−/−^ = 7.76 ± 4.72 APs/burst; *Syt7*^−/−^ = 5.27 ± 3.75 APs/burst; *Doc2α*^−/−^*/Syt7*^−/−^ = 5.36 ± 3.59 APs/burst). WT vs. *Doc2α*^−/−^: *p* = 0.077; WT vs. *Syt7*^−/−^: *p* = 0.030; WT vs. *Doc2α*^−/−^*/Syt7*^−/−^: *p* = 0.144; *Doc2α*^−/−^ vs. *Syt7*^−/−^: \*\*\*\**p* < 0.0001; *Doc2α*^−/−^ vs. *Doc2α*^−/−^*/Syt7*^−/−^: \*\*\*\**p* < 0.0001; *Syt7*^−/−^ vs. *Doc2α*^−/−^*/Syt7*^−/−^: *p* = 0.999. D: No significant change in burst duration in *Doc2α*^−/−^, *Syt7*^*-/*^ and *Doc2α*^−/−^*/Syt7*^−/−^ mice (WT = 268.40 ± 87.42 ms; *Doc2α*^−/−^ = 307.40 ± 75.05 ms; *Syt7*^−/−^ = 244.70 ± 98.97 ms; *Doc2α*^−/−^*/Syt7*^−/−^ = 246.80 ± 99.37 ms). WT vs. *Doc2α*^−/−^: *p* = 0.079; WT vs. *Syt7*^−/−^: *p* = 0.852; WT vs. *Doc2α*^−/−^*/Syt7*^−/−^: *p* = 0.999; *Doc2α*^−/−^ vs. *Syt7*^−/−^: \*\*\**p* = 0.001; *Doc2α*^−/−^ vs. *Doc2α*^−/−^*/Syt7*^−/−^: \*\*\**p* = 0.001; *Syt7*^−/−^ vs. *Doc2α*^−/−^*/Syt7*^−/−^: *p* = 0.999. In panels B, C and D, data were analyzed using the Kruskal-Wallis test with Dunn’s multiple comparisons test. For bursts per minute data, WT = 17 cells from 9 slices from 6 mice; *Doc2α*^−/−^ = 17 cells from 11 slices from 7 mice; *Syt7*^−/−^ = 17 cells from 9 slices from 5 mice; *Doc2α*^−/−^*/Syt7*^−/−^ = 18 cells from 8 slices from 4 mice. For APs/burst and burst duration data, WT = 62 bursts; *Doc2α*^−/−^ = 85 bursts; *Syt7*^−/−^ = 84 bursts; *Doc2α*^−/−^*/Syt7*^−/−^ = 90 bursts. Graphs represent mean ± SD. ns: no significance.

### AR event frequency is unaltered in *Doc2α-* and *Syt7*-KO SC-CA1 synapses

Both Doc2α and Syt7 have been proposed as Ca^2+^ sensors that trigger asynchronous neurotransmitter release (Yao *et al*., 2011; Bacaj *et al*., 2013; Luo & Südhof, 2017; Turecek & Regehr, 2018; Weingarten *et al*., 2024; Wu *et al*., 2024). Moreover, in our previous study (Yao *et al*., 2011), KD of *Doc2α* reduced reverberatory activity in cultured hippocampal preparations. Hence, we tested whether AR is altered in *Doc2α-* or *Syt7-*KO at SC-CA1 synapses (as a proxy for CA3-CA3 synapses). Considering the complex dendritic arborization of CA1 pyramidal neurons, we used slightly larger tip electrodes and recorded eEPSCs with 8-12 MΩ average *R*_s_, which allowed us to quantify small-amplitude distal events occurring far from the soma. Since each CA1 pyramidal neuron receives inputs from numerous Schaffer collaterals, we estimated the frequency of these AR events by normalizing to the evoked synchronous EPSC amplitude in 100 consecutive trials. We did not observe any change in the overall number of AR events (Fig. 4B and C; *p* = 0.702, Kruskal-Wallis test with Dunn’s multiple comparisons test; WT = 0.68 ± 0.17 events/nA; *Doc2α*^−/−^ = 0.74 ± 0.28 events/nA; *Syt7*^−/−^ = 0.75 ± 0.30 events/nA; *Doc2α*^−/−^*/Syt7*^−/−^ = 0.58 ± 0.24 events/nA; WT = 9 cells from 9 slices from 7 mice; *Doc2α*^−/−^ = 8 cells from 8 slices from 5 mice; *Syt7*^−/−^ = 8 cells from 8 slices from 5 mice; *Doc2α*^−/−^*/Syt7*^−/−^ = 8 cells from 7 slices from 4 mice.). Under near-physiological conditions, the AR event frequency peaks around 30-60 ms after the stimulus in SC-CA1 synapses. We note that the observed AR frequency was lower than in other brain regions, but this could be due, in part, to the slightly different experimental conditions used for recording (Turecek & Regehr, 2018; Weingarten *et al*., 2024). Surprisingly, we did observe a significant increase in the quantal amplitude in the Doc2α/Syt7 DKOs, but not in the single KOs (Fig. 4D; *p* = 0.004; Kruskal-Wallis test with Dunn’s multiple comparisons test; (WT = −21.89 ± 1.95 pA; *Doc2α*^−/−^ = −22.52 ± 3.28 pA; *Syt7*^−/−^ = −21.37 ± 1.71 pA; *Doc2α*^−/−^*/Syt7*^−/−^ = −26.62 ± 2.37 pA; WT = 9 cells from 9 slices from 7 mice; *Doc2α*^−/−^ = 8 cells from 8 slices from 5 mice; *Syt7*^−/−^ = 8 cells from 8 slices from 5 mice; *Doc2α*^−/−^*/Syt7*^−/−^ = 8 cells from 7 slices from 4 mice. These findings will require further study. The main finding here is that the observed increase in burst activity in the KOs is not due to changes in AR.

### Intrinsic properties of CA3 pyramidal neurons are unchanged in *Doc2α-* and *Syt7-* KO mice

Since increased burst generation could result from changes in the intrinsic properties of CA3 neurons (Pinsky & Rinzel, 1994; Shao & Dudek, 2009), we examined these parameters via whole-cell current-clamp recordings. We patched neurons in the CA3a/b border, because of their higher recurrent collateral connectivity (Fig. 5A), which plays a role in burst initiation, and because numerous studies suggest that bursts originate in CA3a/b and spread to CA3c and CA1 via Schaffer collaterals (Buzsáki, 2015; Raus Balind *et al*., 2019). For these experiments, spontaneous synaptic activity was suppressed with 100 µM picrotoxin, 10 µM CNQX, and 50 µM APV, and a membrane potential of −70 mV was maintained throughout the recordings. In the first measurement, we found that the RMP (V_*rest*_) of the CA3 pyramidal neurons was unchanged (Fig. 5D; *p* = 0.864, Brown-Forsythe ANOVA test with Dunnett T3 multiple comparisons test; (WT = −70 ± 3.58 mV; *Doc2α*^−/−^ = −69.39 ± 2.73 mV; *Syt7*^−/−^ = −70.29 ± 2.91 mV; *Doc2α*^−/−^*/Syt7*^−/−^ = −70.34 ± 2.99 mV; WT = 20 cells from 11 slices 7 mice; *Doc2α*^−/−^ = 11 cells from 8 slices from 6 mice; *Syt7*^−/−^ = 15 cells from 8 slices from 6 mice; *Doc2α*^−/−^*/Syt7*^−/−^ = 15 cells from 9 slices from 5 mice). We then assessed the firing rate of CA3 neurons by injecting a depolarizing current in 25 pA increments, from 0 to 200 pA. We did not observe any significant changes in the mean firing rate at various current steps (Fig. 5B and C, *p* = 0.833, Brown-Forsythe ANOVA test with Dunnett T3 multiple comparisons test; WT = 17 cells from 9 slices from 7 mice; *Doc2α*^−/−^ = 17 cells from 10 slices from 6 mice; *Syt7*^−/−^ = 16 cells from 9 slices from 6 mice; *Doc2α*^−/−^*/Syt7*^−/−^ = 16 cells from 9 slices from 5 mice). We then examined the input resistance (R_*input*_) of these neurons by injecting hyperpolarizing current in increments of 20 pA, from 0 to −200 pA. The overall R_*input*_ was not significant between the groups (Fig. 5E and F, *p* = 0.0643, Brown-Forsythe ANOVA with Dunnett T3 multiple comparisons test; (WT = 154.50 ± 24.19 MΩ; *Doc2α*^−/−^ = 141 ± 29.82 MΩ; *Syt7*^−/−^ = 156 ± 19.32 MΩ; *Doc2α*^−/−^*/Syt7*^−/−^ = 165 ± 26.57 MΩ; WT = 17 cells from 9 slices from 7 mice; *Doc2α*^−/−^ = 17 cells from 10 slices from 6 mice; *Syt7*^−/−^ = 16 cells from 9 slices from 6 mice; *Doc2α*^−/−^*/Syt7*^−/−^ = 16 cells from 9 slices from 5 mice). Further, the action potential amplitude (Fig. 5G and H, *p* = 0.104, Kruskal-Wallis test with Dunn’s multiple comparisons test; (WT = 118.90 ± 7.28 mV; *Doc2α*^−/−^ = 112.60 ± 8.71 mV; *Syt7*^−/−^ = 110.80 ± 14.82 mV; *Doc2α*^−/−^*/Syt7*^−/−^ = 112 ± 11.51 mV; WT = 17 cells from 9 slices from 7 mice; *Doc2α*^−/−^ = 17 cells from 10 slices from 6 mice; *Syt7*^−/−^ = 16 cells from 9 slices from 6 mice; *Doc2α*^−/−^*/Syt7*^−/−^ = 16 cells from 9 slices from 5 mice) and half-width (Fig. 5G and I, *p* = 0.509, Kruskal-Wallis test with Dunn’s multiple comparisons test; (WT = 0.87 ± 0.21 ms; *Doc2α*^−/−^ = 0.73 ± 0.12 ms; *Syt7*^−/−^ = 0.96 ± 0.28 ms; *Doc2α*^−/−^*/Syt7*^−/−^ = 1.03 ± 0.47 ms; WT = 17 cells from 9 slices from 7 mice; *Doc2α*^−/−^ = 17 cells from 10 slices from 6 mice; Syt7^−/−^ = 16 cells from 9 slices from 6 mice; *Doc2α*^−/−^*/Syt7*^−/−^ = 16 cells from 9 slices from 5 mice). Overall, we did not observe significant changes in the intrinsic properties of CA3 pyramidal neurons, suggesting that the increased burst activity in the KOs could result from alterations in SV or DCV release.

**Figure 4.**
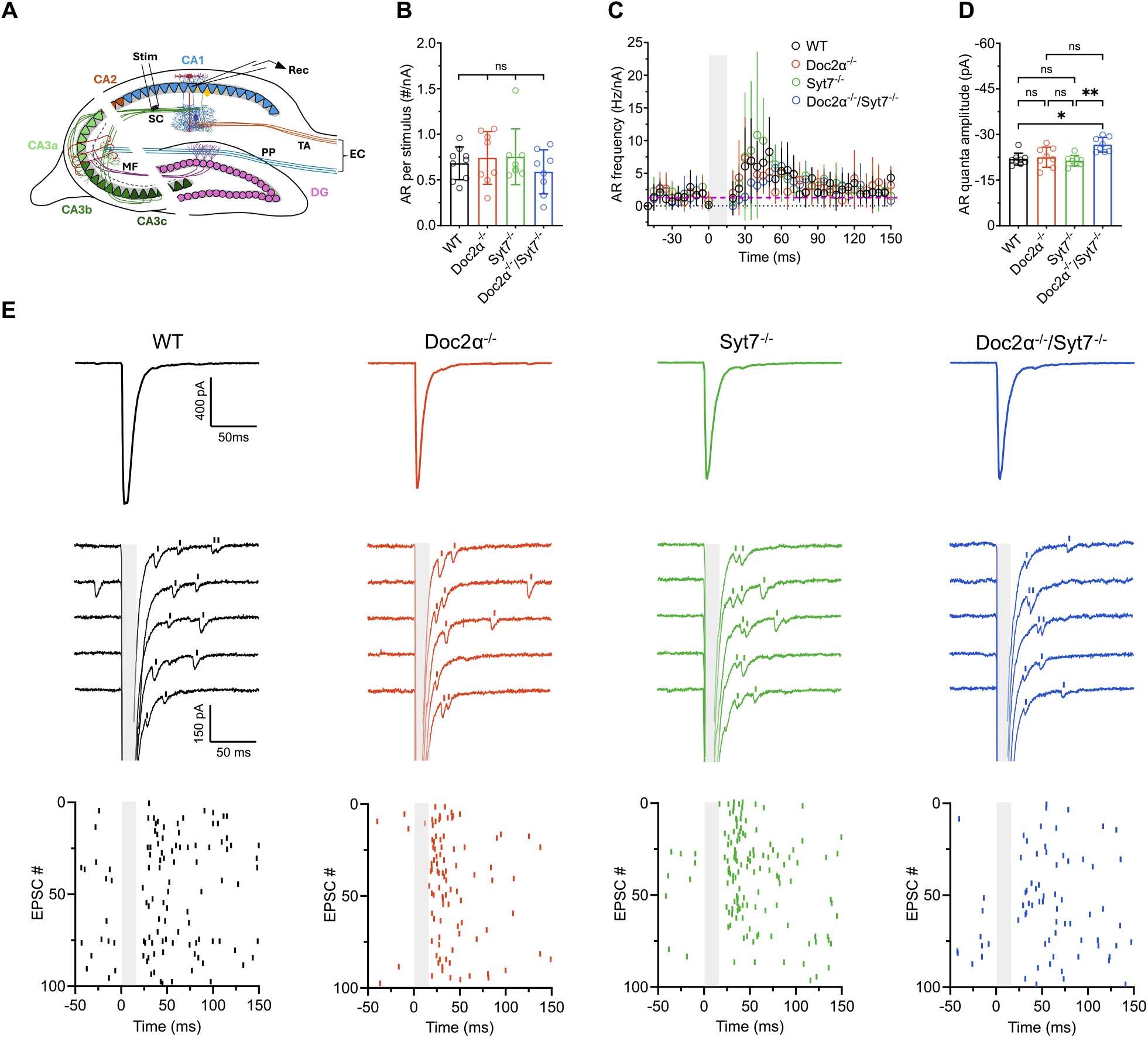
Asynchronous quantal event frequency is unaltered at the Schaffer collateral-CA1 synapses in *Doc2α*^−/−^, *Syt7*^−/−^ *and Doc2α*^−/−^*/Syt7*^−/−^ mice. A: Schematic diagram of the hippocampal slice preparation showing the placement of the stimulating electrode and the whole-cell recording electrode. Schaffer collaterals were severed from the CA3 somas to prevent recurrent activity. B and C: Number of AR events normalized to the synchronous EPSC amplitude (nA) and time course of the normalized AR frequency in 5ms bins, respectively. AR events peaked at 30-60ms after the stimulus. In panel C, the grey shaded area indicates synchronous response, and the dashed purple line represents the network-dependent spontaneous activity. No significant change in the number of AR events/nA between the groups (WT = 0.68 ± 0.17 events/nA; *Doc2α*^−/−^ = 0.74 ± 0.28 events/nA; *Syt7*^−/−^ = 0.75 ± 0.30 events/nA; *Doc2α*^−/−^*/Syt7*^−/−^ = 0.58 ± 0.24 events/nA). WT vs. *Doc2α*^−/−^: *p* = 0.999; WT vs. *Syt7*^−/−^: *p* = 0.999; WT vs. *Doc2α*^−/−^*/Syt7*^−/−^: *p* = 0.999; *Doc2α*^−/−^ vs. *Syt7*^−/−^: *p* = 0.999; *Doc2α*^−/−^ vs. *Doc2α*^−/−^*/Syt7*^−/−^: *p* = 0.999; *Syt7*^−/−^ vs. *Doc2α*^−/−^*/Syt7*^−/−^: *p* = 0.999. D: AR event quantal amplitude is increased in *Doc2α*^−/−^*/Syt7*^−/−^ but not in *Doc2α*^−/−^ and *Syt7*^−/−^ mice (WT = −21.89 ± 1.95 pA; *Doc2α*^−/−^ = −22.52 ± 3.28 pA; Syt7^−/−^ = −21.37 ± 1.71 pA; *Doc2α*^−/−^*/Syt7*^−/−^ = −26.62 ± 2.37 pA). WT vs. *Doc2α*^−/−^: *p* = 0.999; WT vs. *Syt7*^−/−^: *p* = 0.999; WT vs. *Doc2α*^−/−^*/Syt7*^−/−^: *p* = 0.018; *Doc2α*^−/−^ vs. *Syt7*^−/−^: *p* = 0.999; *Doc2α*^−/−^ vs. *Doc2α*^−/−^*/Syt7*^−/−^: *p* = 0.146; *Syt7*^−/−^ vs. *Doc2α*^−/−^*/Syt7*^−/−^: \**p* = 0.006. E: Representative average of 100 synchronous EPSCs elicited by SC stimulation (top panel), representative synaptic responses showing AR events (middle panel) and raster plots of AR events from 100 consecutive trials (bottom panel). AR events in the decay phase of EPSC in the middle panel are highlighted using vertical bars above each raw trace. In panels B and D, data were analyzed using the Kruskal-Wallis test with Dunn’s multiple comparisons test. For AR measurements, WT = 9 cells from 9 slices from 7 mice; *Doc2α*^−/−^ = 8 cells from 8 slices from 5 mice; *Syt7*^−/−^ = 8 cells from 8 slices from 5 mice; *Doc2α*^−/−^*/Syt7*^−/−^ = 8 cells from 7 slices from 4 mice. Graphs represent mean ± SD. ns: no significance; CA: cornu ammonis; DG: dentate gyrus; EC: entorhinal cortex; PP: perforant pathway; MF: mossy fiber pathway; SC: Schaffer collateral pathway; TA: temporoammonic pathway; Stim: stimulating electrode; Rec: recording electrode.

**Figure 5.**
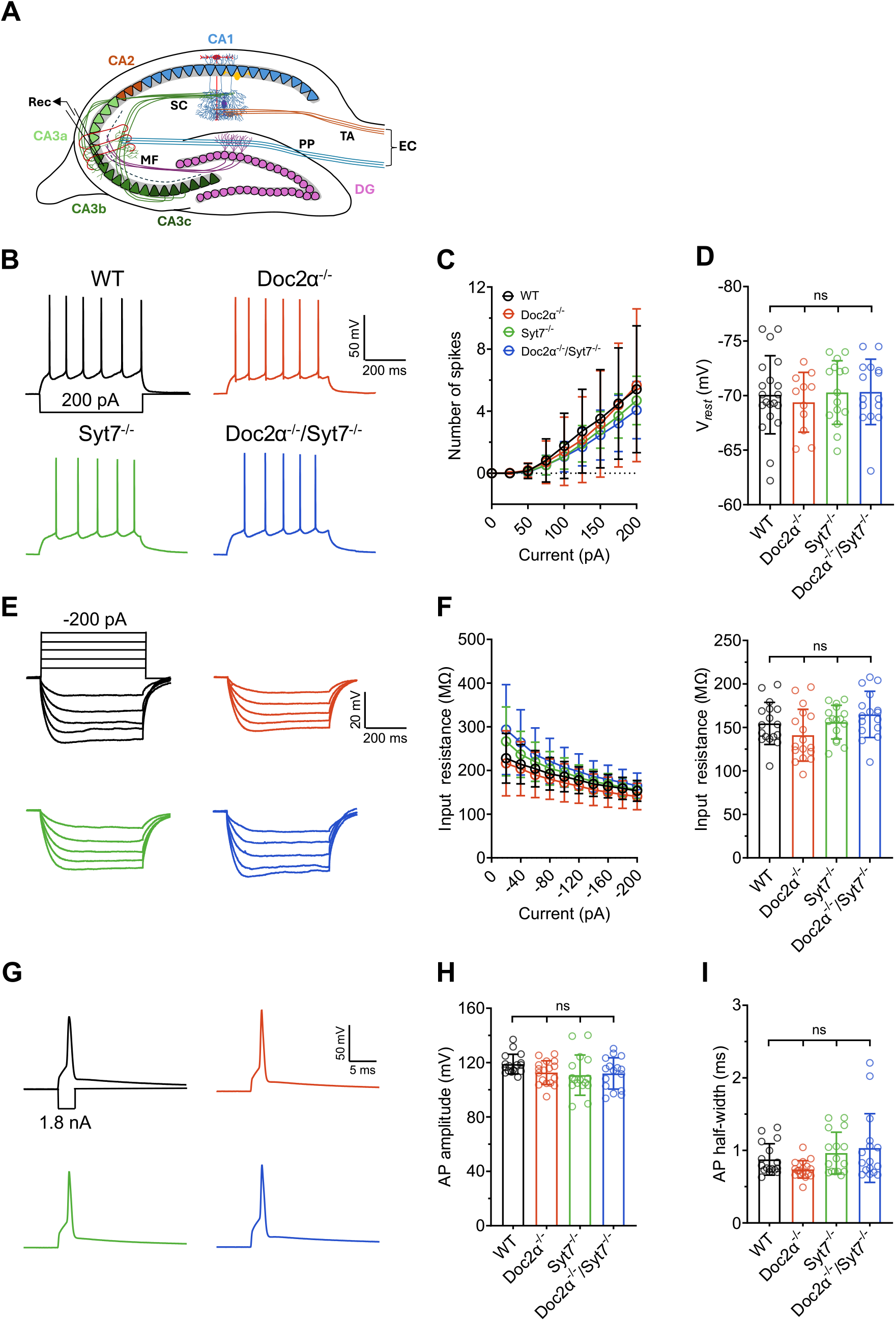
Intrinsic properties of CA3 pyramidal neurons are unaltered in *Doc2α*^−/−^, *Syt7*^−/−^ *and Doc2α*^−/−^*/Syt7*^−/−^ mice. A: Schematic diagram of the hippocampal slice preparation showing the placement of the whole-cell recording electrode in the CA3 subfield. B and C: Representative traces of CA3 neuronal firing in response to 200 pA of depolarizing current, and mean firing rate at various depolarizing current steps, respectively. The firing rates across the various current injection steps did not differ significantly between the groups. WT vs. *Doc2α*^−/−^: *p* = 0.999; WT vs. *Syt7*^−/−^: *p* = 0.996; WT vs. *Doc2α*^−/−^*/Syt7*^−/−^: *p* = 0.973; *Doc2α*^−/−^ vs. *Syt7*^−/−^: *p* = 0.992; *Doc2α*^−/−^ vs. *Doc2α*^−/−^*/Syt7*^−/−^: *p* = 0.957; *Syt7*^−/−^ vs. *Doc2α*^−/−^*/Syt7*^−/−^: *p* = 0.999. D: The resting membrane potential (V_*rest*_) of CA3 neurons was unaltered between the groups (WT = −70 ± 3.58 mV; *Doc2α*^−/−^ = −69.39 ± 2.73 mV; *Syt7*^−/−^ = −70.29 ± 2.91 mV; *Doc2α*^−/−^*/Syt7*^−/−^ = −70.34 ± 2.99 mV). WT vs. *Doc2α*^−/−^: *p* = 0.990; WT vs. *Syt7*^−/−^: *p* = 0.999; WT vs. *Doc2α*^−/−^*/Syt7*^−/−^: *p* = 0.999; *Doc2α*^−/−^ vs. *Syt7*^−/−^: *p* = 0.958; *Doc2α*^−/−^ vs. *Doc2α*^−/−^*/Syt7*^−/−^: *p* = 0.949; *Syt7*^−/−^ vs. *Doc2α*^−/−^*/Syt7*^−/−^: *p* = 0.999. E: Representative traces of hyperpolarized membrane potentials during input resistance (R_input_) measurements at: −40, −80, −120, −160 and −200 pA injected current. F: Input resistance measured at various hyperpolarizing current steps (left panel) and the quantified overall input resistance (right panel). No statistical difference in input resistance measurements (WT = 154.50 ± 24.19 MΩ; *Doc2α*^−/−^ = 141 ± 29.82 MΩ; *Syt7*^−/−^ = 156 ± 19.32 MΩ; *Doc2α*^−/−^*/Syt7*^−/−^ = 165 ± 26.57 MΩ). WT vs. *Doc2α*^−/−^: *p* = 0.619; WT vs. *Syt7*^−/−^: *p* = 0.999; WT vs. *Doc2α*^−/−^*/Syt7*^−/−^: *p* = 0.799; *Doc2α*^−/−^ vs. *Syt7*^−/−^: *p* = 0.430; *Doc2α*^−/−^ vs. *Doc2α*^−/−^*/Syt7*^−/−^: *p* = 0.112; *Syt7*^−/−^ vs. *Doc2α*^−/−^*/Syt7*^−/−^: *p* = 0.850. G: Representative traces of a single action potential elicited by injecting 1.8 nA of depolarizing current. H: Quantified action potential amplitude measured from the baseline to peak. No statistical difference was observed in AP amplitude (WT = 118.90 ± 7.28 mV; *Doc2α*^−/−^ = 112.60 ± 8.71 mV; *Syt7*^−/−^ = 110.80 ± 14.82 mV; *Doc2α*^−/−^*/Syt7*^−/−^ = 112 ± 11.51 mV). WT vs. *Doc2α*^−/−^: *p* = 0.568; WT vs. *Syt7*^−/−^: *p* = 0.116; WT vs. *Doc2α*^−/−^*/Syt7*^−/−^: *p* = 0.454; *Doc2α*^−/−^ vs. *Syt7*^−/−^: *p* = 0.999; *Doc2α*^−/−^ vs. *Doc2α*^−/−^*/Syt7*^−/−^: *p* = 0.999; *Syt7*^−/−^ vs. *Doc2α*^−/−^*/Syt7*^−/−^: *p* = 0.999. I: Quantified action potential half-width data measured from baseline to 50% peak amplitude. No statistical difference was observed in AP half-width (WT = 0.87 ± 0.21 ms; *Doc2α*^−/−^ = 0.73 ± 0.12 ms; *Syt7*^−/−^ = 0.96 ± 0.28 ms; *Doc2α*^−/−^*/Syt7*^−/−^ = 1.03 ± 0.47 ms). WT vs. *Doc2α*^−/−^: *p* = 0.349; WT vs. *Syt7*^−/−^: *p* = 0.999; WT vs. *Doc2α*^−/−^*/Syt7*^−/−^: *p* = 0.999; *Doc2α*^−/−^ vs. *Syt7*^−/−^: *p* = 0.116; *Doc2α*^−/−^ vs. *Doc2α*^−/−^*/Syt7*^−/−^: *p* = 0.132; *Syt7*^−/−^ vs. *Doc2α*^−/−^*/Syt7*^−/−^: *p* = 0.999. In panels C, D and F, data were analyzed using the Brown-Forsythe ANOVA test with Dunnett T3 multiple comparisons test. In H and I, data were analyzed using the Kruskal-Wallis test with Dunn’s multiple comparisons test. For RMP measurements, WT = 20 cells from 11 slices 7 mice; *Doc2α*^−/−^ = 11 cells from 8 slices from 6 mice; *Syt7*^−/−^ = 15 cells from 8 slices from 6 mice; *Doc2α*^−/−^*/Syt7*^−/−^ = 15 cells from 9 slices from 5 mice. For firing rate, input resistance and action potential kinetics measurements, WT = 17 cells from 9 slices from 7 mice; *Doc2α*^−/−^ = 17 cells from 10 slices from 6 mice; *Syt7*^−/−^ = 16 cells from 9 slices from 6 mice; *Doc2α*^−/−^*/Syt7*^−/−^ = 16 cells from 9 slices from 5 mice. Graphs represent mean ± SD. ns: no significance; CA: cornu ammonis; DG: dentate gyrus; EC: entorhinal cortex; PP: perforant pathway; MF: mossy fiber pathway; SC: Schaffer collateral pathway; TA: temporoammonic pathway; Rec: recording electrode.

## DISCUSSION

In this study, we used KO mice to probe the potential functions of Doc2α and Syt7 in network-dependent spontaneous activity in the hippocampus. We found that loss of either of these presynaptic proteins increases burst generation in the hippocampal CA3 subfield in disinhibited slices from juvenile mice (P15-21) without a change in AR. The frequencies and amplitudes of sEPSCs and sIPSCs remain unchanged in the KO mice. Further, our whole-cell current-clamp measurements show that the observed phenotype is not due to changes in the intrinsic properties of CA3 pyramidal neurons, suggesting alterations in SV or DCV release. To the best of our knowledge, this is the first study to report the effect of *Doc2α-* and *Syt7-*KO on burst generation in the hippocampal CA3 subfield.

### The CA3 subfield and burst generation

The hippocampal CA3 subfield is a critical region for associative learning and memory (Joo & Frank, 2018). There are three main inputs to the CA3: i) projections from the entorhinal cortex (perforant pathway), ii) projections from dentate granule cells (mossy fiber pathway), and iii) associative-commissural projections due to RCs (Fig. 1A). All three inputs are spatially segregated into different layers of the CA3 subfields - a, b and c - with CA3a and CA3c being distal and proximal to the dentate gyrus (DG), respectively. CA3a and CA3b axons ramify extensively and form RCs to generate hypersynchronous bursts, a critical physiological function for information transfer and memory consolidation (Buzsáki, 2015; Rebola *et al*., 2017). CA3 burst generation is a complex phenomenon that requires tightly controlled spatiotemporal integration. These bursts are generated by intrinsic mechanisms, network activity, or a combination of both (Zeldenrust *et al*., 2018). A plethora of cellular and molecular mechanisms have been proposed to modulate this synaptic integration (Buzsáki, 2015). Recent evidence suggests that CA3 neurons generate bursts due to Ca^2+^ spikes that produce slow afterdepolarization in the soma by prolonging dendritic plateau potentials (Raus Balind *et al*., 2019). These plateau potentials play a role in the induction of behavioral time-scale plasticity (BTSP) in the CA3 (Li *et al*., 2024; Madar *et al*., 2025; Magee, 2026). In addition, imbalances in burst generation can lead to pathological epileptiform activity as seen in neurological or neurodevelopmental disorders (Goldberg & Coulter, 2013; Buzsáki, 2015).

### CA3 burst activity is increased in *Doc2α-* and *Syt7-*KO hippocampi independent of alterations in synaptic transmission

We reiterate that Doc2 proteins act as Ca^2+^ sensors that regulate the spontaneous release of SVs, also called miniature neurotransmitter release or minis, in acute slices and cultured hippocampal neurons; their relative contribution to minis in the hippocampus depends on the specific synapse under study (Groffen *et al*., 2010; Ramirez *et al*., 2017; Courtney *et al*., 2018; Khan & Regehr, 2020; Wang *et al*., 2023). However, loss of Doc2α-triggered minis is unlikely to regulate bursting activity, since paired recordings showed that the RCs of CA3 neurons have unitary EPSPs of ~0.56 mV, which is insufficient to trigger an action potential in the postsynaptic cell (Guzman *et al*., 2016). In addition, there is no clear evidence for a role of Syt7 in the regulation of minis in wild-type neurons (Chen *et al*., 2017; Luo & Südhof, 2017). These findings argue against a role for minis in the bursting behavior described in the current study.

In the *Doc2α*- and *Syt7-*KOs, we found no change in the asynchronously released quanta at SC-CA1 synapses following a single stimulus. These findings contrast previous results, in which we observed reduced AR in the *Doc2α-*KO, whereas disruption of *Syt7* had no effect on AR triggered by a single stimulus (Wu *et al*., 2024). These differences might be due to differences in experimental conditions, where, in the past, we used higher external calcium (2 mM Ca^2+^) in the aCSF recorded from 6-8 week old adult mice. Indeed, AR is a common mechanism that is supported by both Doc2α and Syt7 at a variety of other synapses (Chen *et al*., 2017; Turecek & Regehr, 2018; Deng *et al*., 2020; Wu *et al*., 2024). Loss of AR at an excitatory-excitatory synapse, e.g. the RCs in CA3, should result in reduced excitation. However, a previous study demonstrated prolonged bursts, but decreased burst initiation probability and onset rate, in the CA3 subfield after substituting Ca^2+^ with Sr^2+^ in aCSF (Jones *et al*., 2007). Sr^2+^ is slowly cleared from the cytosol, so it enhances AR (Xu-Friedman & Regehr, 2000; Turecek & Regehr, 2018; Weingarten *et al*., 2024). Interestingly, we observed an increased burst phenotype in CA3 without any changes in AR quanta under near-physiological conditions (i.e., 1.2 mM Ca^2+^ and 33-34 °C), where AR is typically low. Our results have a caveat in that we measured the AR from CA3 terminals at the SC-CA1 synapse rather than at the recurrent synapses. The CA3 RCs make it difficult to quantify AR with the traditional field-stimulation approach because of recurrent network activity, so this was not further explored. Taken together, it is unlikely that changes in the slow, asynchronous mode of transmission underlies the observed changes in CA3 burst activity, but this cannot be ruled out within the CA3 RCs. Optical methods, such as iGluSnFR-based approaches, might make it possible to overcome technical limitations and enable analysis of AR within the RCs of the CA3 region, but at present, this approach suffers from limited temporal resolution.

### An alternate hypothesis

In addition to neurotransmitter release, both Doc2α and Syt7 regulate peptide release from DCVs in neuroendocrine cells (MacDougall *et al*., 2018). Neuropeptides secreted by the inputs to CA3 regulate burst firing of pyramidal neurons. Namely, neuropeptide-Y (NPY) and C-type natriuretic peptide (CNP) suppress the generation of SPWs in the CA3 subfield by decreasing feed-forward excitatory transmission (Klapstein & Colmers, 1993; Decker *et al*., 2009; Pollali & Draguhn, 2025). The CA3 subfield layers, *stratum radiatum* and *stratum oriens*, are enriched in NPY receptors (Klapstein & Colmers, 1993). NPY is secreted by DG and CA3 interneurons that often co-express somatostatin (Pelkey *et al*., 2017). NPY suppresses glutamate release by reducing Ca^2+^ influx in nerve terminals (Colmers *et al*., 1988). It is important to note that NPY does not alter the passive or active membrane properties of CA3 neurons.

NPY is of particular interest since mossy fiber terminals, the major inputs to the CA3, are also rich in NPY-bearing DCVs (Henze *et al*., 2000). However, a recent study showed that the selective knockout of Syt7, induced via AAV injection in the DG (to disrupt expression in mossy fibers), reduced the likelihood of CA3 population activity *in vivo* by reducing spike transfer from DG to CA3 (Marneffe *et al*., 2026), which would seem to be inconsistent with our observations. Rather, Marneffe et al. (2026) proposed that abrogation of facilitation was the underlying mechanism. However, the authors estimate a ~60% loss of Syt7 in dentate granule cells - due to incomplete coverage of the virally induced knockout - with DG and CA3 interneurons spared. So, it remains possible that loss of Syt7 disrupts the release of burst-suppressing neuropeptides such as NPY, and thereby promotes burst activity.

Extending this hypothesis to Doc2 proteins, we reiterate that, at the mRNA level, dentate granule cells express both *Doc2α* and *Doc2β*, whereas CA3 cells express only *Doc2α* (Verhage *et al*., 1997), but it is not known if both isoforms are translated, and isoform-specific antibodies are not currently available. We did not observe the burst phenotype in *Doc2β* KOs (data not shown), so either Doc2β is not expressed, or it fails to regulate a neuropeptide that controls bursting, or our hypothesis regarding mossy fiber control of bursting activity is incorrect. In addition, whether Doc2α regulates neuropeptide release from neurons has been called into question in studies using overexpressed NPY fusion proteins in cultured hippocampal neurons (van Westen *et al*., 2021). This will require careful re-examination, as Doc2α has again been shown to regulate the DCVs release in neuroendocrine and non-neuronal cells (Friedrich *et al*., 2008; Ramalingam *et al*., 2012; Houy *et al*., 2017; MacDougall *et al*., 2018).

While there are arguments for and against the inhibitory neuropeptide hypothesis, we reiterate that this remains a leading model, since the only remaining untested function that is conserved between Doc2α and Syt7 - albeit across different cell types - is the control of DCV release. In this light, it is notable that the bursting phenotype was not additive in the CA3 subfield from *Doc2α* and *Syt7* double KO mice. This finding is consistent with the idea that both proteins act in the same pathway. Further testing of the DCV hypothesis will require measurements of neuropeptide release within the CA3 region of *Doc2α-* and *Syt7-* knockout mice, perhaps via development of G-protein-coupled activation-based (GRABs) sensor for NPY and other candidate peptides.

### Limitations of the study

The current study utilized juvenile mice, so it is possible that the phenotype we observed is limited to this developmental period. In this light, we note that Sholl analysis of dentate granule cells showed increased neurite branching in *Doc2α*-KO mice (Wang *et al*., 2023). Increased branching of axons and dendrites during postnatal development is one mechanism underlying CA3 hyperexcitability (Gómez-Di Cesare *et al*., 1997). Notably, the observed increase in branching complexity was absent in adult mice. We also did not characterize the behavioral phenotypes exhibited by the *Doc2α-* and *Syt7-*KO mice. Although both the KOs are viable and appear normal, recent reports suggest they may have developmental issues (Shen *et al*., 2020; Wang *et al*., 2023). For example, loss of Doc2α has been proposed to cause autism spectrum-like disorder in mice, in the same age group as the mice used in our study (Wang *et al*., 2023). Finally, it is not yet known whether there is an increase in SPW generation in *Doc2α-* and *Syt7-*KO mice *in vivo*, or whether the increased burst activity interferes with memory-dependent tasks. These are important areas of inquiry in future studies.

In conclusion, using juvenile *Doc2α-* and *Syt7-*KO mice, we uncover a novel burst-generating phenotype in the hippocampal CA3 subfield. Our study has implications for understanding the roles of these presynaptic proteins in regulating excitatory transmission in hippocampal circuits and their associated physiological functions.

## GENERATIVE AI STATEMENT

No generative AI was used in planning, executing the experiments and writing the manuscript.

## AUTHORS’ CONTRIBUTION

Conceptualization: RJS and ERC; Investigation: RJS; Data curation and analysis: RJS; Funding acquisition: ERC; Visualization: RJS and ERC; Writing: RJS and ERC.

## DATA AND MATERIALS AVAILABILITY

All the raw data is available on Dryad, a publicly accessible repository. https://doi.org/10.5061/dryad.wpzgmsc50

## ACKNOWLEDGMENTS

We thank Dr. Dennis Weingarten and Dr. Skyler Jackman for their suggestions and for helping with AR experiments. We thank Divya Teja Pasam for writing Excel macros to automate AR data analysis. We thank Dr. Ion Razvan Popescu, a senior scientist in the lab, for providing specific input and critical suggestions, and all other members of the Chapman lab for helpful discussions and support. ERC is an Investigator of the Howard Hughes Medical Institute (HHMI). We also thank the SciDraw artists Georgia Papadimitriou and Federico Claudi for providing the illustrations of the hippocampus (https://doi.org/10.5281/zenodo.14367775) and the pyramidal neuron (https://doi.org/10.5281/zenodo.3925905) for the graphical abstract, respectively. The illustrations are licensed under CC-BY 4.0.

